# Phylogeographic analysis of *Begomovirus* coat and replication-associated proteins

**DOI:** 10.1101/2023.11.08.565637

**Authors:** Alvin Crespo-Bellido, J. Steen Hoyer, Yeissette Burgos-Amengual, Siobain Duffy

## Abstract

Begomoviruses are globally distributed plant pathogens that significantly limit crop production. These viruses are traditionally described according to phylogeographic distribution and categorized into two groups: begomoviruses from the Africa, Asia, Europe, and Oceania (AAEO) region and begomoviruses from the Americas. Monopartite begomoviruses are more common in the AAEO region while bipartite viruses predominate in the Americas, where the begomoviruses lack the V2/AV2 gene involved in inter-cellular movement and RNA silencing suppression found in AAEO begomoviruses. While these features are generally accepted as lineage-defining, the number of known species has doubled due to sequence-based discovery since 2010. To reevaluate the geographic groupings after the rapid expansion of the genus, we conducted phylogenetic analyses for begomovirus species representatives of the two longest and most conserved begomovirus proteins: the coat and replication-associated proteins. Both proteins still largely support the broad AAEO and Americas begomovirus groupings, except for sweetpotato-infecting begomoviruses that form an independent, well-supported clade for their coat protein regardless of the region they were isolated from. Our analyses do not support more fine-scaled phylogeographic groupings. Monopartite and bipartite genome organizations are broadly interchanged throughout the phylogenies and the absence of the V2/AV2 gene is highly reflective of the split between Americas and AAEO begomoviruses. We observe significant evidence of recombination within the Americas and within the AAEO region, but rarely between the regions. We speculate that increased globalization of agricultural trade, the invasion of polyphagous whitefly vector biotypes and recombination will blur begomovirus phylogeographic delineations in the future.

## INTRODUCTION

The *Geminiviridae* family consists of plant-infecting viruses with broad host range that can severely constrain agricultural crop production [1]. Genus *Begomovirus* constitutes the most speciose out of the 14 currently approved geminivirus genera, with 445 established species listed in the current Master Species List of the International Committee on the Taxonomy of Viruses (MSL38 v2, https://ictv.global/msl). The begomoviruses are whitefly-transmitted, mostly dicot-infecting pathogens that cause significant crop losses in tropical and subtropical regions around the world [2]. The success of begomoviruses as emerging pathogens is facilitated by international agricultural trade, the widespread distribution of the polyphagous whitefly vector (*Bemisia tabaci* cryptic species complex), and adaptation fueled by high mutation frequencies and frequent genetic exchange through recombination and reassortment [3–5].

Historically, two major begomovirus groups have been recognized based on phylogeographic distribution and evolutionary features: begomoviruses originally identified in Africa, Asia, Europe, and Oceania (AAEO) region, and begomoviruses first isolated from the Americas [6, 7]. These “AAEO” and “Americas” begomovirus designations are used in lieu of the traditional phrases “Old World” and “New World” for accuracy throughout this work, given that Australia – which is lumped with Afro-Eurasia for begomovirus classification – is generally considered part of “the New World” (including in the wine industry). While widely accepted, not all begomovirus species follow this coarse geographic grouping. The sweetpotato-infecting begomoviruses (known as “sweepoviruses”) are not split at continent scales, but rather form a distinct clade, with some viruses discovered in samples collected in the Americas and others discovered in samples from the AAEO region [4, 8, 9]. An additional group of legume-infecting begomoviruses known as the “legumoviruses” is also considered a distinct lineage from the two broad regional groups, although all legumoviruses have been sampled in the AAEO region [8, 10, 11].

All begomovirus genomes are organized into one (monopartite) or two (bipartite) circular single-stranded DNA (ssDNA) segments, each independently encapsulated in twinned, quasi-icosahedral particles that are characteristic of geminiviruses [11]. Segments range from ∼2.5-2.8kb. For bipartite begomoviruses the two segments are referred to as DNA-A and DNA-B. Monopartite genomes are homologous to the DNA-A segment of bipartite begomoviruses. All infectious genomes described thus far have at least five protein-encoding open reading frames (ORFs). On the virion-sense strand, they possess the *V1* (in monopartites) or *AV1* (in bipartites) gene which codes for the coat protein (CP). The complementary-sense strand encodes the replication-associated protein Rep (*C1/AC1*), the transcriptional activator protein TrAP (*C2/AC2*), a replication enhancer REn (*C3/AC3*), and an RNA-silencing suppressor C4/AC4 (functions reviewed in [12, 13]). The AAEO begomoviruses possess an additional ORF called *V2/AV2* which codes for a protein associated with movement and silencing-suppression [6, 13, 14]. Recent studies have suggested possible functional roles for additional DNA-A ORFs: *C5/AC5* [15–17], *C6* [18], *C7* [19] and *V3* [20, 21]. However, transcription and translation of these ORFs has only been studied in a very small number of begomoviruses, such as the intensively studied, pandemic tomato yellow leaf curl virus (TYLCV) [21].

The DNA-B of bipartite begomoviruses encodes a nuclear shuttle protein NSP in the virion-sense (*BV1*) and an intercellular movement protein MP in the complementary sense (*BC1*). For DNA-B, a recent study showed that an additional ORF *BV2* is translated during infection [22]. The segments of bipartite begomoviruses share a conserved region (CR) of ∼200 nucleotides that includes a stem-loop structure with a conserved nonanucleotide where rolling-circle replication is initiated. Additionally, the CR contains several regulatory elements, including multiple copies of cis elements known as iterons that are specific binding sites for Rep during replication [23].

There is evidence that several features distinguish most begomoviruses isolated in the Americas from their AAEO region counterparts: a different number and arrangement of iterons in the CR [23, 24], a conserved PWRsMaGT motif in the N-terminal domain of CP [14], an RFATDKS motif in the REn protein [25], shorter genome segments [26] and the aforementioned lack of V2/AV2. Additionally, Americas begomoviruses were thought to be exclusively bipartite until recent reports of monopartite begomoviruses in the Americas with Americas-type features [27–32]. It has been suggested that the preponderance of bipartite begomoviruses in the Americas may stem from the lack of V2/AV2 [8, 14], because the V2 protein is thought to be the main mediator of inter-cellular movement during infection for monopartite viruses [16, 33, 34] – although this remains controversial [35]. Under this hypothesis, the absence of the movement capabilities of V2/AV2 increases the reliance of DNA-A on the movement functions of DNA-B proteins and leads to stronger interdependence between the segments [26].

More than 100 species have been discovered since the last large-scale phylogenetic analyses of the genus [8, 9]. As a result, we sought to update our understanding of begomovirus evolution by creating updated, comprehensive phylogenies for the two main proteins in the monopartite/DNA-A genome segments: CP and Rep. We pursued an amino-acid-sequence-based approach instead of evaluating the evolution of the entire monopartite genome/DNA-A segment because extensive genomic divergence results in an unreliable nucleotide alignment, especially in the intergenic regions. We believe that this protein-based approach is phylogenetically robust given that the CP and Rep are the two most conserved begomovirus proteins [7, 12, 36–38] and together they are encoded by ∼70% of nucleotides in the segment. We mapped genomic features and geographic location onto our phylogenies and showed that the evolution of these proteins is largely reflective of the traditional geographic delineations, though there are notable exceptions. The CP and Rep phylogenies are significantly incongruent with each other, suggesting extensive recombination within but not between the two main geographic regions. The vast majority of AAEO begomoviruses are monopartite while most Americas begomoviruses are considered bipartite, although five independent evolutions of a monopartite genome organization in Americas begomoviruses are observed. The results also reveal that the presence/absence of V2/AV2 is highly correlated with CP and Rep evolution and support the absence of V2/AV2 as a lineage-defining feature of Americas begomoviruses. Additionally, our results confirm that genome/DNA-A segment length correlates with presence/absence of V2/AV2.

## METHODS

### Sequence retrieval of begomovirus RefSeq exemplar CP and Rep sequences

Annotated begomovirus coding sequences corresponding to each begomovirus species exemplar with a RefSeq accession number listed in the ICTV Virus Metadata Resource (VMR #18, 2021-10-19, https://ictv.global/vmr) were downloaded from GenBank in protein FASTA file format (June 2023). CP and Rep amino acid sequences were extracted and split into separate data sets for analysis. Several RefSeq sequences were misannotated in NCBI, most commonly with CP and V2/AV2 protein ORFs mislabeled as the other (sequences listed in Supplemental file 1). We confirmed the identity of the ORF products by performing a BLAST search [39] (https://blast.ncbi.nlm.nih.gov/Blast.cgi) of the non-redundant protein database at NCBI and included the homologous sequences in their corresponding data sets. For exemplar sequences missing ORF annotations (listed in Supplemental file 1), ORFfinder (https://www.ncbi.nlm.nih.gov/orffinder/) was used to identify CP and Rep ORFs that were subsequently translated and added to each corresponding data set after BLAST confirmation [39].

Metadata associated with each exemplar in our data set – including country of isolation, geographic designation (i.e., AAEO/Americas), genome segmentation (i.e., monopartite/bipartite), presence/absence of V2/AV2 and length of genome/DNA-A segments – are included in Supplemental file 1. Based on historical context within begomovirology [7], AAEO begomoviruses in this study are defined as those whose exemplars were sampled outside of the Americas whereas Americas begomoviruses are those sampled within North America (including Central America and the Caribbean) and South America. AAEO begomovirus genomes are labeled as bipartite if they are associated in GenBank with a DNA-B segment and monopartite if they are not. Since Americas begomoviruses are generally considered bipartite due to a perceived reliance on a DNA-B segment to successfully establish a systemic infection, exemplars are labeled as bipartite if they are associated in the VMR with a DNA-B segment, monopartite if there is experimental evidence demonstrating their ability to systematically infect the host species from which they were isolated in the absence of a DNA-B segment or ‘undetermined’ if no DNA-B segment had been sampled along with it. The presence/absence of V2/AV2 was determined for each exemplar using ORFfinder and BLAST [39].

### Alignments and phylogenetic analysis

Multiple sequence alignments were constructed using the MUSCLE method [40] as implemented in MEGA 11 [41] and manually corrected using AliView v1.26 [42]. After an initial alignment inspection, exemplars with either severely truncated (i.e., length < 50% of the average length of the protein) or very divergent (i.e., causing us to doubt protein homology) CP or Rep sequences were excluded from the data set (Supplemental table 1). Our final CP and Rep data sets contained amino acid sequences from 432 begomovirus species. Due to the difficulties in aligning the Rep sequences at the N- and C-terminal ends, the alignment was trimmed to eliminate all residues prior to the iteron related domain (i.e., the known Rep functional region closest to the Rep start [23]) in the N-terminus and after a conserved geminivirus motif found near the C-terminus, which corresponds to where other circular, Rep-encoding single-stranded DNA viruses possess an arginine finger motif [43, 44]. Alignments have been archived as Zenodo records at https://zenodo.org/record/8338685.

Maximum likelihood (ML) trees were inferred with IQ-Tree v2.0.7 [45] using the best fitting substitution model identified by the built-in ModelFinder feature [46]. Tree inference was performed with 3000 ultrafast bootstrap (UFBoot) replicates, a perturbation strength of 0.2 and a stopping rule requiring an iteration interval of 500 iterations between unsuccessful improvements to the local optimum. The -bnni flag was enabled to reduce the risk of overestimating branch supports with UFBoot due to severe model violations. Midpoint-rooted consensus phylogenies were annotated and visualized using Treeviewer [47]. Tree files in NEXUS format available on Zenodo at https://zenodo.org/record/8338685. [48–50]

### Begomovirus exemplar genome/DNA-A length distributions

The distribution of monopartite genome and bipartite DNA-A segment lengths for each begomovirus exemplar was calculated using R version 4.2.0 [51] and visualized using the *gghistogram* function in the *ggpubr* R package [52].

### Shannon entropy scores for CP and Rep data sets

Shannon entropy scores-which describe the amount of information within a variable [53] (in this case, amino acid sequence alignments)– were calculated as a measure of protein diversity for sets of aligned protein sequences using an adapted Python script from [54] available at https://github.com/acrespo-virevol/shannon-entropy.

### Comparison of begomovirus exemplar CP and Rep phylogenies

A tanglegram of the CP and Rep ML phylogenies was created using the *dendextend* R package [55]. The ML trees were input into R and converted into dendrograms for the tanglegram analysis using the *ape* R package [56]. The tanglegram was disentangled by iteratively rotating inner branches of one tree while the other remained static until the entanglement could not be reduced any further using the ‘step2side’ method with *dendextend*.

## RESULTS

The 432 begomovirus species representatives in our data set were sampled across 59 countries and classified as either Africa-Asia-Europe-Oceania (AAEO) or Americas begomoviruses (Figure 1). There is unevenness in the data set with respect to sampling region, with 253 sequence exemplars sampled in the AAEO region and 179 sequence exemplars sampled in the Americas. India (n=65), Brazil (n=60) and China (n=53) are the three countries with the most sequence exemplars in the data set, with no other country providing more than 30 species representatives (Figure 1).

**Figure 1.**
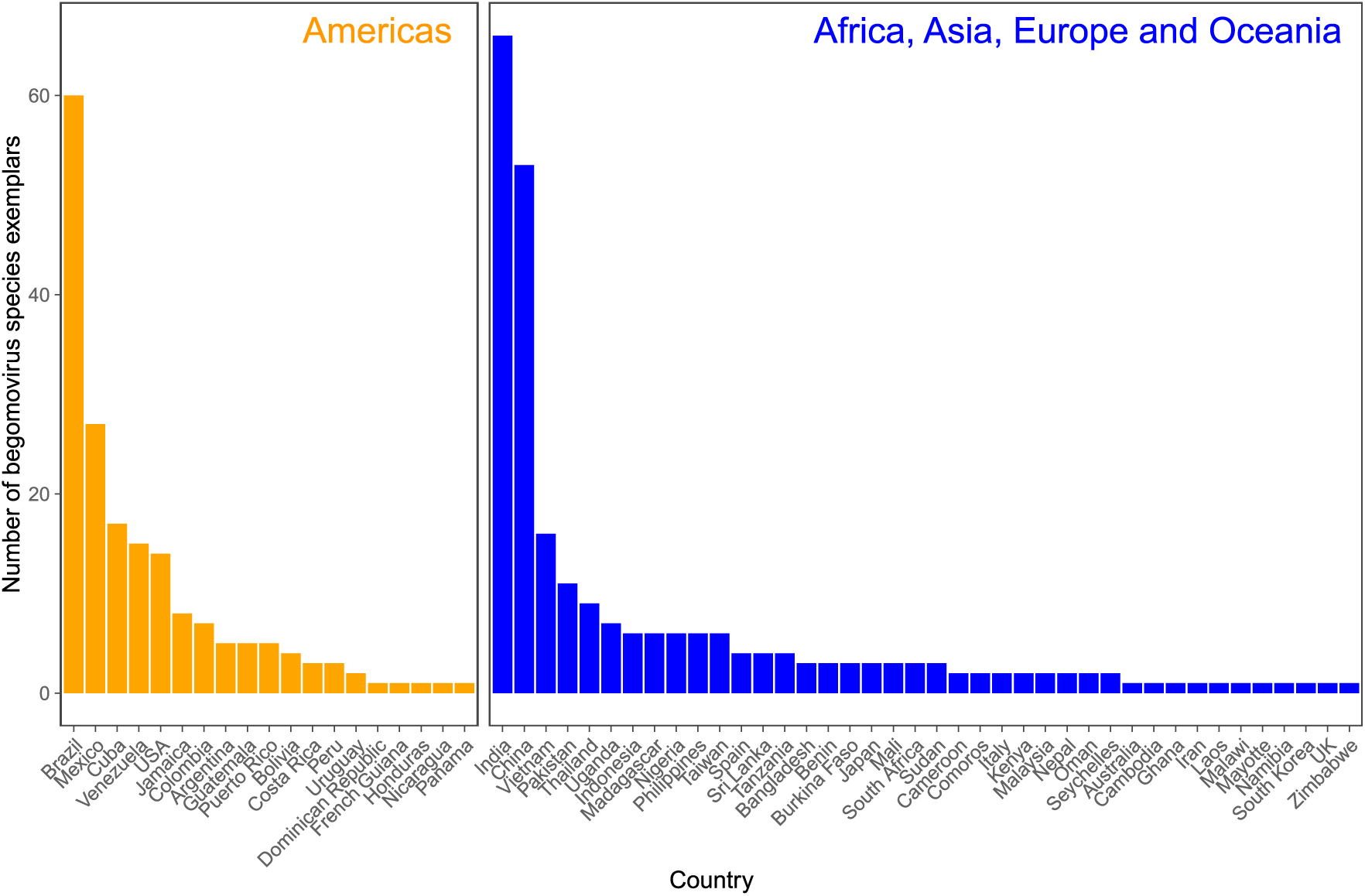
Number of begomovirus species exemplars in this study (n= 432) by country of isolation. Begomovirus exemplar counts are divided into Africa, Asia, Europe, and Oceania (AAEO) and Americas groups based on the country where the viruses were sampled.

### The Americas begomoviruses, which lack V2/AV2, are distinct

The maximum likelihood (ML) phylogeny for the complete CP amino acid sequence alignment is shown in Figure 2. The midpoint root separates the sweepoviruses from the rest of the sequences, indicating that sweepovirus CPs - which were sampled in Asia, Europe, and North and South America - have significantly diverged from those of all other begomoviruses (distance from the midpoint root depicted in Fig. S1). Aside from the sweepoviruses, the phylogenetic analysis supports an Americas-type CP monophyletic clade with the sole exception of the tomato latent virus (ToLV) exemplar from Cuba, a known interspecific recombinant that inherited its CP from the AAEO TYLCV introduced to the Americas in the 1990s [57, 58]. On the other hand, the Corchorus yellow vein virus (CoYVV) exemplar from Vietnam is the sole begomovirus sampled in the AAEO region for which the CP clusters with Americas sequences. This virus and Corchorus golden mosaic virus (CoGMV), which was also identified in Vietnam [14, 59], have been described as a distinct AAEO begomovirus lineage with Americas begomovirus-type features that is basal to the Americas begomovirus clade based on phylogenetic analysis of DNA-A and DNA-B segments [8, 14]. In our analysis, the CoGMV CP does appear to be basal to the Americas clade (Fig. 2). However, the CoYVV CP is closely related to sequences deeply nested within the Americas clade, which suggests that the CoYVV CP originates from a contemporaneous Americas-type lineage.

**Figure 2.**
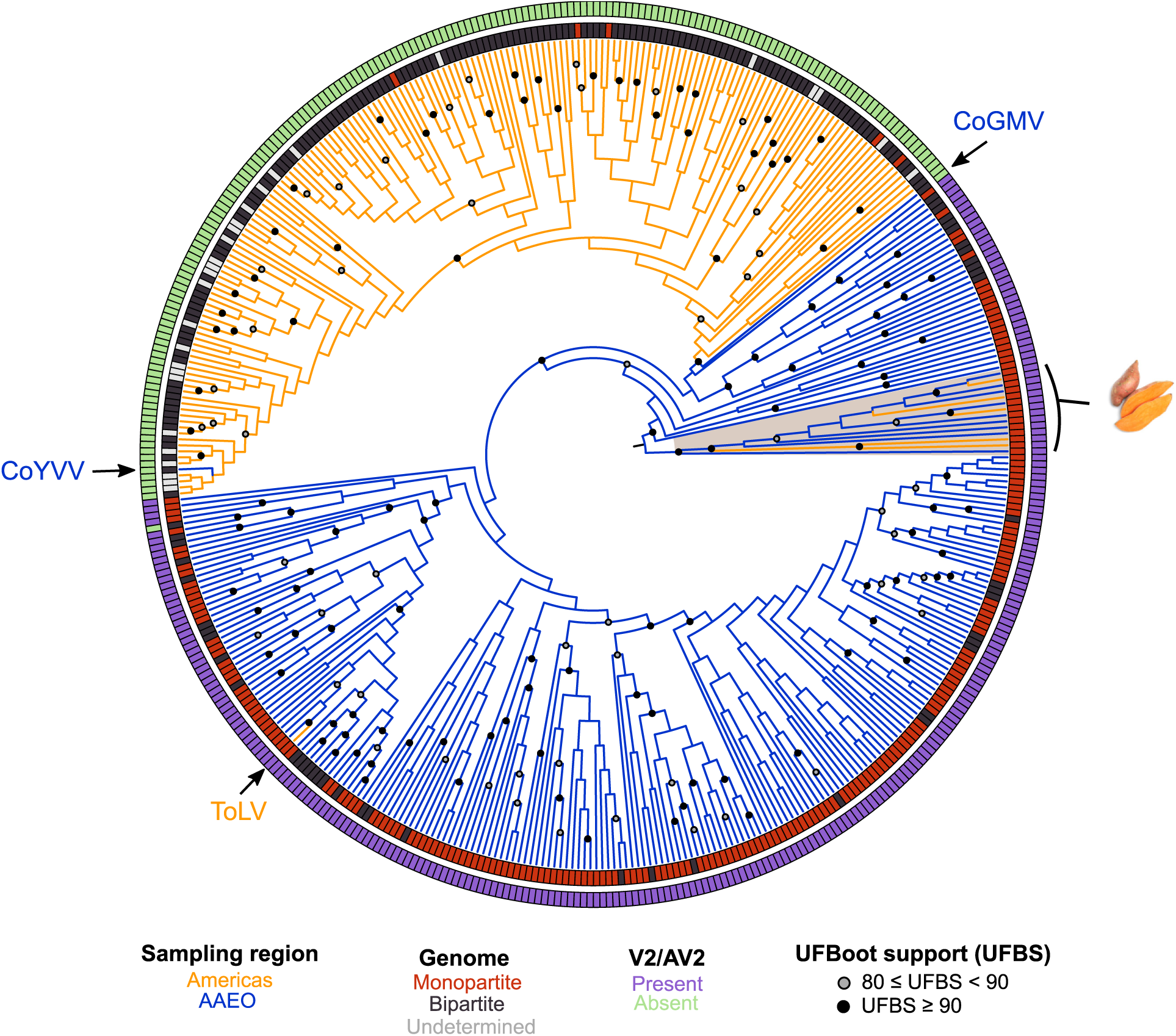
Circular midpoint-rooted maximum likelihood phylogenetic tree of complete CP amino acid sequences of 432 begomovirus RefSeq species exemplars. The maximum likelihood phylogenetic tree was constructed using IQ-Tree v2.07 with automatic selection of the best-fit substitution model (JTT+I+G4). Tree inference was performed with 3000 ultrafast bootstrap (UFBoot) replicates and a stopping rule of 500 iterations between unsuccessful improvements to the local optimum. Only UFboot branch support values ≥ 80% are shown. Branches are colored based on the region where the exemplar was sampled - Americas exemplars in orange and AAEO exemplars in blue. Sweetpotato-infecting viruses are highlighted and denoted by an image of sweetpotato (public domain). CoGMV= Corchorus golden mosaic virus (RefSeq accession number NC_009644), CoYVV= Corchorus yellow vein virus (RefSeq accession number NC_006358), ToLV= tomato latent virus (RefSeq accession number NC_038963).

Outside of the sweepovirus clade (whose members are all monopartite), we see that most of the AAEO sequences are classified as monopartite (194/242=80%) whereas most of the Americas exemplars are confirmed to be bipartite (142/176=81%, Fig. 2; Fig. S1). The CP phylogeny does not support a single origin for bipartitism from a monopartite ancestor, nor a single origin of monopartism from bipartitism in the Americas. There are five monopartite exemplars from the Americas in our data set with demonstrated ability to systematically infect wild host species (disregarding the highly susceptible experimental lab host *Nicotiana benthamiana*) in the absence of a DNA-B segment - Corchorus yellow vein Cuba virus [32], tomato leaf curl purple vein virus [30], tomato leaf deformation virus [28, 60], tomato mottle leaf curl virus [27] and tomato twisted leaf virus [29] - and their placement on the tree indicates that their monopartite genome structure has evolved multiple times via independent loss of DNA-B.

The presence/absence of V2/AV2 is more reflective of the evolution of the begomovirus CP than geographic region or genome segmentation (Fig. 2, Fig. S1). We observe that every exemplar in the Americas-type CP clade lacks a V2/AV2 whereas all other exemplars except the bipartite soybean chlorotic blotch virus (SbCBV) possess a V2/AV2 gene. The absence of V2/AV2 is supported as a clade-defining feature of begomoviruses in the Americas. Given the fact that the SbCBV CP is closely related to other CPs of begomoviruses with an V2/AV2 gene – which is consistent with analyses of the entire DNA-A segment [61] - it is likely that the lack of an AV2 homolog represents a recent deletion of AV2 in SbCBV.

Further analysis of the sequences reveals that clades are not structured by continent (Fig. S1). The sweepovirus CP clade is strongly supported even though exemplars were sampled across four continents. The Americas-type CPs do not show a separation between North and South American exemplars and are significantly intermixed. Interestingly, 46/51 African sequences cluster together in a well-supported clade (highlighted in Fig. S1). This mostly-African clade includes the CP of a Spanish representative of the highly invasive and recombinogenic tomato yellow leaf curl virus [62, 63] and two of its descendants through recombination – ToLV from Cuba [57] and tomato leaf curl Liwa virus from Oman [64]. It also includes a close relative of TYLCV, tomato yellow leaf curl Sardinia virus (TYLCSaV) from Italy, along with two descendent lineages – tomato yellow leaf curl Axarquia virus and tomato yellow leaf curl Malaga virus, both from Spain – that stemmed from separate recombination events with TYLCV [65, 66]. The close relationship of the European TYLCV and TYLCSaV sequences with African exemplars may suggest an African origin for these tomato-infecting viruses. An additional Asian exemplar from Oman within the clade, okra leaf curl Oman virus, is also a recombinant that inherited its CP from the African cotton leaf curl Gezira virus [67]. The last member of the clade is the whitefly-associated begomovirus 7 from Spain, which has yet to be characterized in detail but is a close relative of African tomato-infecting viruses based on analysis of the DNA-A [68].

### The begomovirus Rep phylogeny is incongruent with the CP phylogeny

The ML phylogeny for the trimmed Rep amino acid sequence alignment (see Methods) is shown in Figure 3. The midpoint root separates the Americas from the AAEO and sweepovirus exemplars. In contrast to the CP phylogeny, the sweepovirus Rep sequences are not highly diverged and cluster within the AAEO clade. Interestingly, the Rep sequences of sweepoviruses do not constitute a monophyletic clade, as the sweetpotato leaf curl Hubei virus (SPLCHbV) Rep falls outside of the main sweepovirus cluster (Fig. 3, Fig. S2). It is probable that the SPLCHbV Rep was inherited through recombination with a distantly related AAEO virus. In the Rep phylogeny, ToLV clusters within the Americas clade with its closest relative, the bipartite Sida golden mottle virus (SiGMoV), which is consistent with previous recombination analyses identifying a SiGMoV-like virus as the likely parent for the ToLV Rep [57]. The CoGMV and CoYVV Rep sequences are closely related to each other and to other contemporaneous AAEO sequences. Except for the sweepoviruses, the Rep phylogeny supports AAEO and Americas begomovirus Reps as separate monophyletic clades.

**Figure 3.**
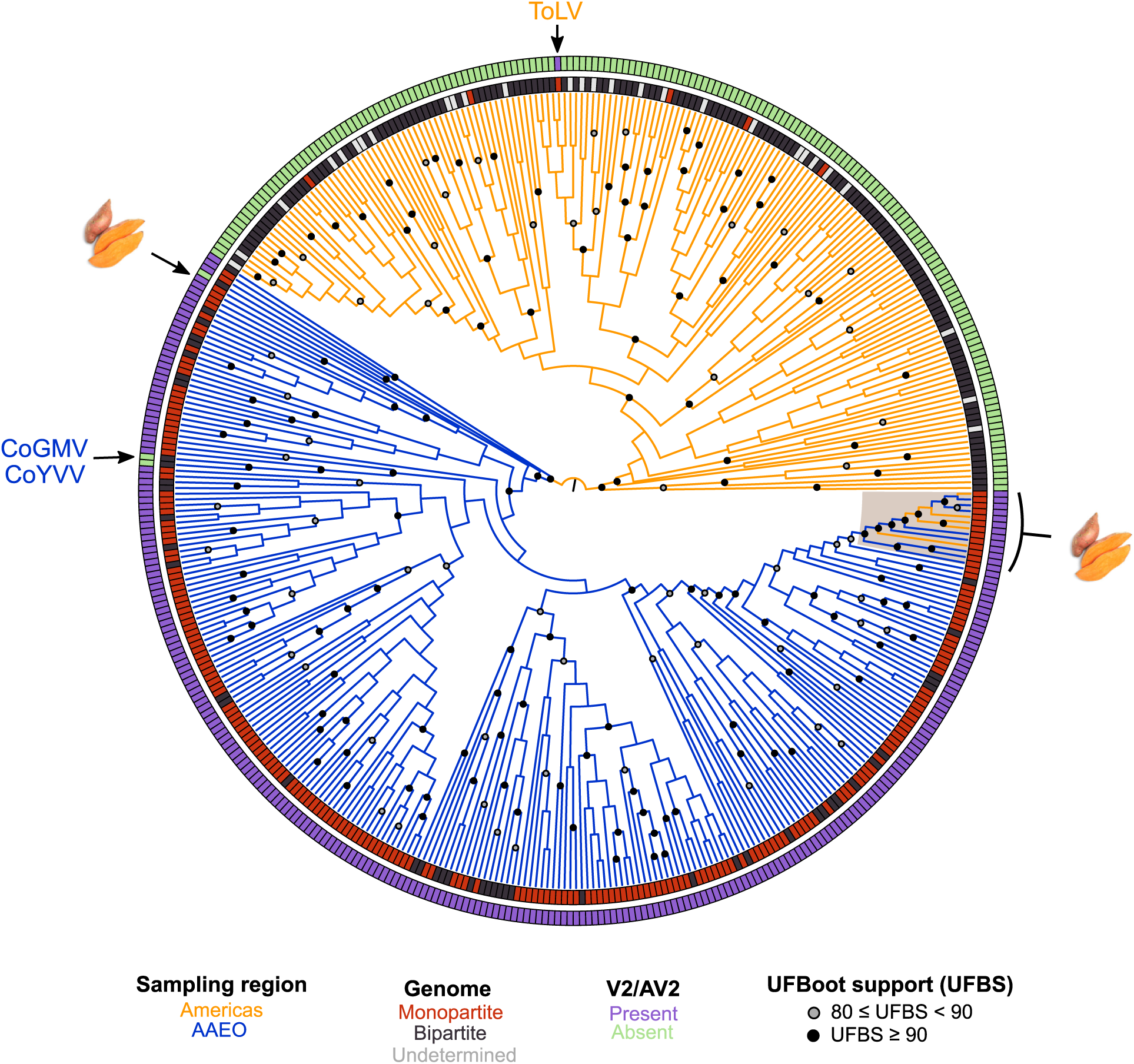
Circular midpoint-rooted maximum likelihood phylogenetic tree of trimmed Rep amino acid sequences of 432 begomovirus RefSeq species exemplars. The maximum likelihood phylogenetic tree was constructed using IQ-Tree v2.07 with automatic selection of the best-fit substitution model (LG+I+G4). Tree inference was performed with 3000 ultrafast bootstrap (UFBoot) replicates and a stopping rule of 500 iterations between unsuccessful improvements to the local optimum. Only UFboot branch support values ≥ 80% are shown. Branches are colored based on the region where the exemplar was sampled - Americas exemplars in orange and AAEO exemplars in blue. Sweetpotato-infecting viruses are highlighted and denoted by an image of sweetpotato (public domain). CoGMV= Corchorus golden mosaic virus (RefSeq accession number NC_009644), CoYVV= Corchorus yellow vein virus (RefSeq accession number NC_006358), ToLV= tomato latent virus (RefSeq accession number NC_038963).

As in the CP phylogeny, the Rep tree does not support a single origin for bipartite genomes from a monopartite ancestor. Additionally, the five monopartite begomoviruses from the Americas do not appear to be closely related based on the evolutionary relationships of their Reps. ToLV presents an interesting case in that its major parent is likely a bipartite begomovirus that acquired the CP and V2 through recombination [57], which resulted in a shift towards a monopartite genome organization.

The Rep phylogeny does not show clustering at the continent level and the well-supported African clade in the CP is not observed for Rep (Fig. S2). Interestingly, the TYLCV and TYLCSaV Reps also cluster with African sequences although the clade is not well-supported.

A comparison of the CP and Rep trees reveals a significant level of incongruency between the phylogenies (Fig. 4), which is suggestive of extensive recombination. The tanglegram suggests high levels of recombination within the Americas and AAEO regions but few cases between them. The exceptions are exemplars already discussed: ToLV, CoYVV and CoGMV. The sweepoviruses also display high levels of incongruency between protein phylogenies, likely related to the large divergence between sweepovirus CP and other CP sequences.

**Figure 4.**
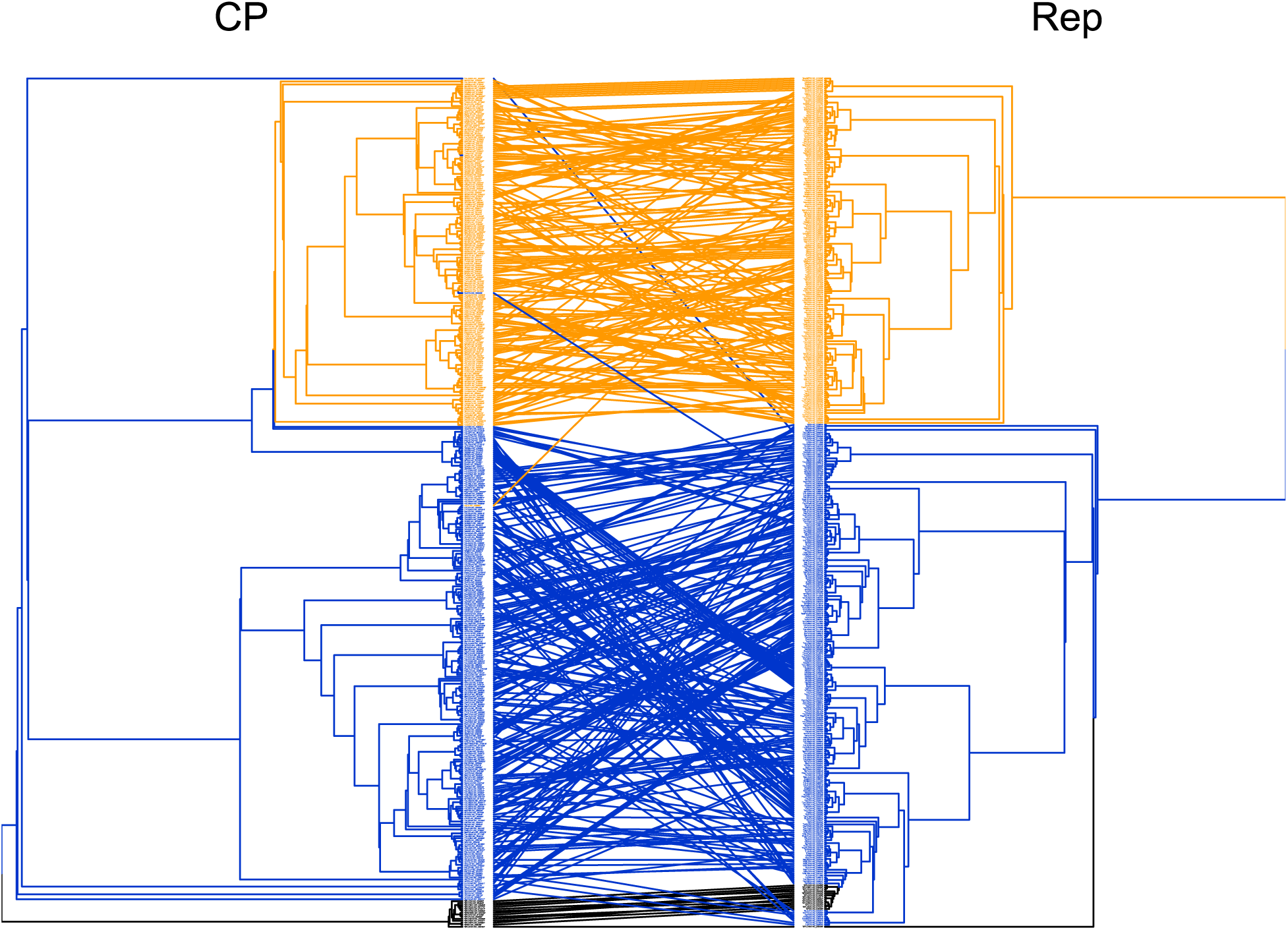
Tanglegram of CP and Rep phylogenies for 432 begomovirus RefSeq species exemplars. Sequences corresponding to the same exemplar are connected by colored lines between each phylogeny. Exemplars from the Americas are orange, AAEO sequences in blue and sweepoviruses are in black.

### More diversity in the CP and Rep of AAEO begomoviruses compared to Americas begomoviruses

We estimated Shannon entropy scores as a general measure of protein diversity for each alignment. Since the CP sequences of the sweepoviruses are so diverged, we removed them from both the CP and Rep data sets when assessing diversity for the AAEO and Americas data sets. We placed the ToLV CP in the AAEO data set and the CoYVV and CoGMV CPs in the Americas-type data set given that the ToLV CP clusters with AAEO sequences while CoYVV and CoGMV are more closely related to Americas CP sequences (Fig. 2). The Shannon entropy scores are higher for AAEO sequences (CP= 0.448; Rep= 0.506) than for Americas sequences (CP= 0.192; Rep= 0.376), indicating higher overall diversity in AAEO begomoviruses.

### Genome/DNA-A length is correlated with geographic region and the presence/absence of V2/AV2

It has been suggested that the absence of V2/AV2 in Americas begomoviruses may be a consequence of one or multiple deletions totaling >100 nucleotides (nt) of its proximal promoter region [26]. We decided to explore the distribution of monopartite genome/bipartite DNA-A segment lengths in the context of AAEO, Americas and sweepovirus groupings, and presence/absence of the V2/AV2 ORF (Fig. 5). The lengths of genome/DNA-A segments largely follow a regional distribution, with the more-frequently-bipartite begomoviruses from the Americas having, on average, segments ∼130 nt shorter (mean length= 2626.5 nt) than the AAEO region viruses (mean length= 2754.6 nt). The monopartite sweepoviruses, regardless of location, have among the largest genomes (mean length= 2792.9 nt). The strong correlation between geographic region and genome/DNA-A segment length appears to be largely due to the presence (mean length= 2757.3 nt) and absence (mean length= 2627.1 nt) of V2/AV2.

**Figure 5.**
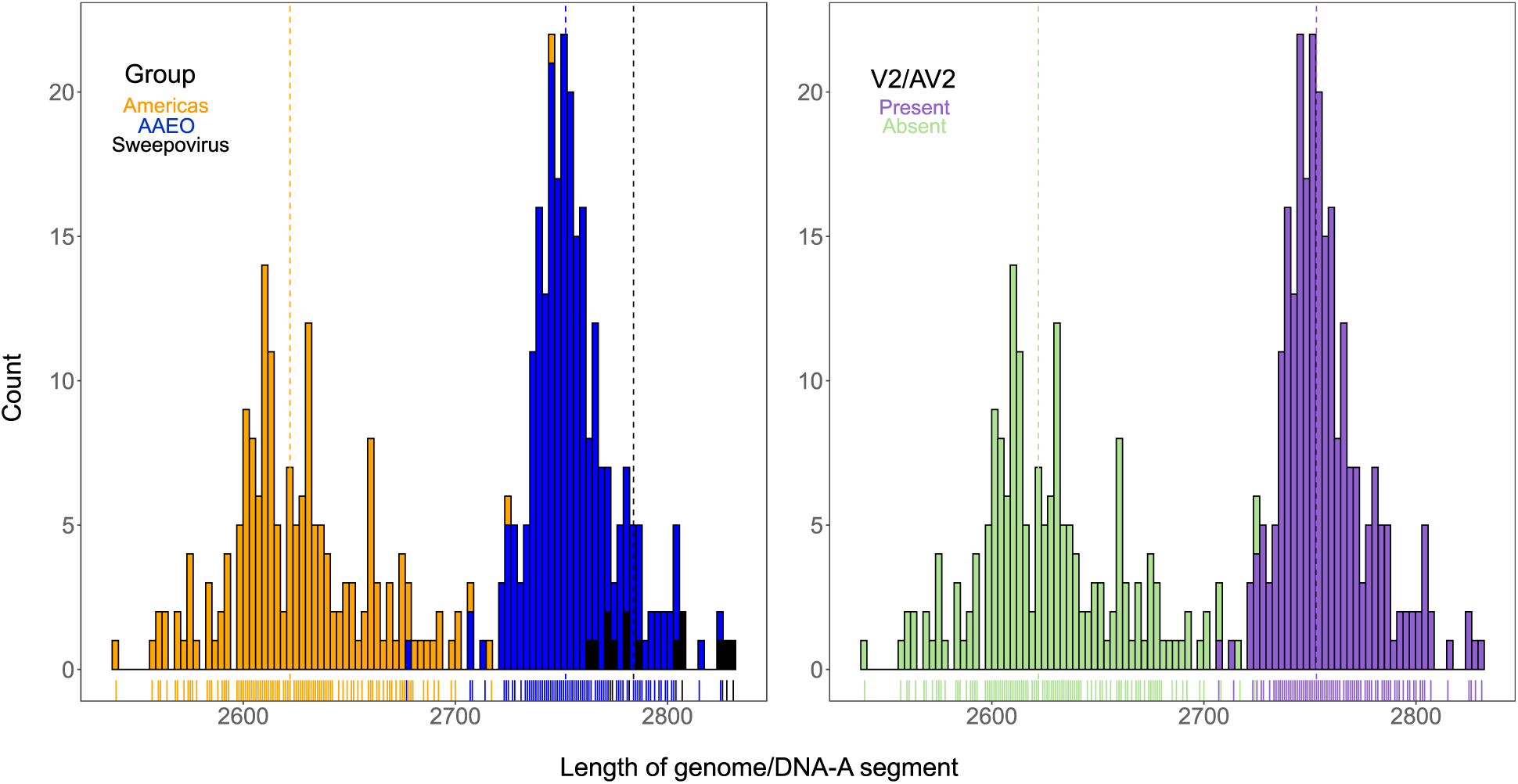
Genome/DNA-A length distribution for three begomovirus groups: Americas, AAEO and sweepoviruses (left) and genome/DNA-A length distribution for begomovirus exemplars by presence/absence of V2/AV2 (right). The length of genomic segments is in number of nucleotides (nt). Mean values for each group are denoted by colored, dashed vertical lines. Americas mean length= 2626.5 nt; AAEO mean length= 2754.6 nt; sweepovirus mean length= 2792.9 nt; absent V2/AV2 mean length= 2627.1 nt; present V2/AV2 mean length= 2757.3 nt.

## DISCUSSION

The *Begomovirus* genus represents an economically important group of emergent plant pathogens that has significantly expanded in the last decade through sequencing-based discovery [5]. We sought to construct robust and updated phylogenies that will serve as resources for future comparative analyses. Using one exemplar sequence per virus species has obvious limitations (particularly for inferences about host range) but was effective for our purpose here. We confirmed the major groupings proposed by Briddon et al. [8], supported by Mondal et al. [9], and used in the 9^th^ ICTV report [10]: sweepoviruses are distinct from other begomoviruses, and the two distinct groups of viruses from AAEO and the Americas. We do not see support for breaking down the geography within the regions more finely – and even the more specifically named clades in the previous analysis had representatives from non-titular countries [8]. We also do not observe an equivalent to the “outsiders” group highlighted by Briddon et al. in either phylogeny.

We observe well-supported clades that include legume-infecting begomoviruses from Asia in both of our analyses (Fig. S1 and S2), which correspond to a previously defined lineage of begomoviruses classified as the “legumoviruses” (also known as “legume yellow mosaic viruses”) that includes mungbean yellow mosaic virus, mungbean yellow mosaic India virus, Dolichos yellow mosaic virus, horsegram yellow mosaic virus and kudzu mosaic virus [8, 69] along with more recently characterized close relatives from Asia: velvet bean severe mosaic virus [70], Rhynchosia yellow mosaic virus [71] and Rhynchosia yellow mosaic India virus [72]. Two other legume-infecting begomoviruses from Africa referenced as “legumoviruses” in the literature – soybean mild mottle virus and Desmodium mottle virus [61] – cluster with the Asian “legumoviruses” for CP but not for Rep. Two additional African viruses referred to as “legumoviruses” – soybean chlorotic blotch virus and cowpea golden mosaic virus – cluster together in a separate clade for both CP and Rep. The evolutionary relationships and overall polyphyly for these African and Asian legume-infecting begomoviruses agrees with previous analyses based on DNA-A segments [61] but conflicts with a more recent analysis claiming monophyly for the African and the Asian species [73]. Interestingly, other legume-infecting exemplars from Asia – such as French bean leaf curl virus, senna leaf curl virus and pea leaf distortion virus – are not closely related to the African/Asian “legumovirus” clade. As legume-infecting begomoviruses are more broadly distributed, we suggest that researchers should minimize use of the word “legumovirus” going forward.

The CP phylogeny supports three designations: a monophyletic, V2/AV2-less Americas clade (including CoYVV and CoGMV), a paraphyletic AAEO group and a sweepovirus clade. On the other hand, the Rep phylogeny is largely consistent with two groups: an Americas clade and AAEO group (including CoYVV, CoGMV and sweepoviruses). However, the lack of V2/AV2 is much more reflective than geography of the evolution of CP and Rep, with SbCBV as the sole exception for CP and the recombinant ToLV and putative recombinants CoYVV and CoGMV as the exceptions for Rep.

### Begomoviruses remain largely delimited by geography

The evolutionary split between the AAEO and Americas begomoviruses was proposed to be a consequence of the dissolution of the Bering land bridge between Asia and North America 20-30 MYA, which may have significantly limited dispersal between these regions until more recent human migration [74]. Increasing globalization of agricultural trade has been linked with plant virus emergence and epidemics, stemming from encounters between introduced crops and the native viruses of a region, climate change, and the spread of plant viruses and their vectors throughout the world [75–77]. There are a few examples of begomovirus introductions across regions such as TYLCV throughout the Americas [58, 78, 79], Abutilon mosaic virus in the United Kingdom [38] and New Zealand [80], squash leaf curl virus in the Middle East [81] and watermelon chlorotic stunt virus in North America [82, 83]. As a result, viral exchange between the AAEO and Americas regions could be reflected in these phylogenies. Past introductions may also be revealed by the presence of cross-region hybrid species, such as ToLV in the Americas and potentially CoYVV and CoGMV in the AAEO region. Yet, we see no other examples of sequences clustering outside the region they were sampled from. It is possible that introduced viral species have not been recovered due to uneven and insufficient sampling; although the detection of novel begomovirus species has more than doubled in the last decade, sequencing is concentrated in certain countries (Fig. 1) which may be biasing the recovered diversity of begomoviruses. Sequencing also has largely been focused on economically important crops, an approach that underestimates the diversity that exists in wild plants and weeds that act as viral reservoirs and have been implicated as sources for recombinant lineages and novel virus species [84–87]. Additionally, while high-throughput sequencing methods are established as essential tools for plant virus discovery [88, 89], infrastructure gaps still hinder plant virus discovery and management in some regions of the world [90]. As sequencing continues to become broader and more accessible, more introduced viral species may be discovered.

Vector-virus co-adaptation might also contribute to the lack of cross-region begomovirus species spread. The CP interacts with many proteins from the vector (and its endosymbionts) in ways that determine transmission and transmission efficiency [91–93]. It is possible that local whitefly biotypes are not well-adapted to support the efficient transmission of introduced begomoviruses, limiting their potential for spread. This could be one reason for the largely-African clade in the CP tree (but not the Rep tree): there may be an undetermined molecular evolutionary signature of adaptation to African whitefly populations that unite these sequences. However, we may expect localized co-adaptation to break down in the future with the introduction of invasive vector biotypes or species between regions. For instance, the introduction of the invasive Middle East-Asia Minor 1 (MEAM1, formerly known as B) and Mediterranean (MED, formerly known as Q) whitefly biotypes in the Americas may facilitate the emergence of introduced AAEO begomoviruses and of new recombinant species. MEAM1 exhibits greater polyphagy and can colonize tomato more efficiently than native populations, which has resulted in tomato epidemics across South America through the transference of indigenous begomoviruses from wild plant hosts into tomato [94–96]. The greater host range, along with increased capacity for environmental adaptation, may also explain the gradual displacement of local whitefly populations by MEAM1 in most agricultural regions of Latin America [94, 97–99]. The MED biotype is also a successful invader in the Americas [94, 100–102], and is resistant to pesticides compared to MEAM1 and indigenous whiteflies [103, 104], so it will likely also be successful at displacing indigenous whiteflies in managed ecosystems. Ultimately, the continuous spread of these invasive vector species increases the likelihood that a co-adapted, introduced AAEO begomovirus will successfully spread and be detected. It may also promote coinfection in a wider variety of hosts and, consequently, the conditions for the emergence of novel recombinants. The relationship between different whiteflies and monopartite and bipartite viruses are just beginning to be elucidated [105–107], and future work may connect the success of monopartite begomoviruses in the Americas with cross-region, introduced whitefly vectors.

Recombination is a major mechanism of speciation for begomoviruses [4, 108, 109]. Although we see frequent recombination within regions, cross-region recombinants are rare in our analyses (Fig. 5). AAEO-Americas begomovirus recombination events may be under strong negative selection, given that recombination between significantly diverged viruses could disrupt intragenomic co-adaptation and selection will act to maintain co-evolved protein-protein and protein-DNA interactions [110]. However, the few examples of emergent cross-region recombinants (ToLV and CoYVV) demonstrate that it is possible. Without significant improvements in quarantine and more careful trade of planting materials, it can be anticipated that more begomoviruses will be introduced from AAEO to the Americas and vice versa, increasing the chances for inter-region recombination.[110–114]

### V2/AV2 function likely varies across the genus

Both the CP and Rep trees (Fig 2, 3) show that monopartite and bipartite genome organizations are frequently interchanged in the AAEO region (although monopartite viruses appear much more prevalent) and that monopartite viruses evolved at least five separate times from bipartite relatives in the Americas. It has been suggested that the V2/AV2 of AAEO bipartite begomoviruses is a vestigial gene from monopartite ancestors that may increase fitness but is no longer required for movement [115], and that the loss of the V2/AV2 gene led to the prevalence of bipartite viruses in the Americas [8, 14]. This conventional wisdom has broken down with additional sampling and experimentation. For instance, Sri Lankan cassava mosaic virus is a bipartite virus, but its DNA-A alone can infect *Nicotiana benthamiana* (although coinfecting with its DNA-B increases symptom development) [116]. A similar pattern has been found with two other AAEO bipartite viruses: tomato yellow leaf curl Thailand virus in tomato [117] and tomato leaf curl Gujarat virus in *N. benthamiana* [118]. Corresponding results have been seen with in *N. benthamiana* with the DNA-A of two Americas begomoviruses (tomato chlorotic mottle virus [119] and Sida golden mosaic Braco virus [120]), and the fact that these infectious clones were unable to replicate in the solanaceous hosts in which they were found may imply that they need a DNA-B for successful infection of some hosts. That some bipartite DNA-As can infect some hosts without DNA-B, and that infectious clones of DNA-A segments are successful in the hyper-susceptible laboratory host *N. benthamiana* [121] but not originally identified field hosts make the designations of monopartite and bipartite more fluid than expected. Similarly, the confirmed monopartite pepper yellow vein Mali virus is frequently associated with a DNA-B component in the field, and laboratory testing showed that it infects more cells, replicates to higher DNA levels and transmits more effectively when coinfecting with a DNA-B [122]. That begomoviruses might be functionally monopartite and bipartite in different hosts, or under different conditions, helps explain frequent transitions between monopartite and bipartite genome organizations in the CP and Rep phylogenies.

The intra- and inter-cellular movement proteins on the DNA-B segment may not be as essential in all hosts as previously assumed – they could lend a competitive advantage against other viruses in some hosts, and/or be needed to counter a host defense pathway in others. Conversely, the movement functions of the DNA-B proteins may be adequately performed in the monopartite viruses (and the DNA-A segments capable of infecting as independent infectious clones) by other proteins. The CP of AAEO monopartite begomoviruses, aided by host factors and other viral proteins like V2 and C5, possesses the nucleocytoplasmic trafficking functions for intracellular movement that are carried out by the nuclear-shuttle protein in the DNA-B segment of bipartite begomoviruses [34, 123, 124]. Indeed, the nuclear shuttle protein may be a distant homolog of the CP [125]. Cell-to-cell movement is attributed mainly to V2 in monopartite viruses [34, 115, 124, 126], with assistance of C5 for docking to the plasmodesmata [16]. Yet, the multiple examples of monopartite Americas begomoviruses lacking V2 [27–32, 60] suggest that V2 is not essential for systemic movement for all begomoviruses. It is possible that other proteins may supply the movement functions of V2 in its absence, as begomovirus proteins are highly multifunctional [13, 127]. For example, a study with the TYLCV-Israel strain showed that loss-of-function V2 mutants retain the ability to establish systemic yet attenuated infection [35]. However, abolishing the expression of C4 – which is already implicated in movement in some species – inhibited systemic infection [35]. Additionally, a newly discovered small gene, V3, can target plasmodesmata and partly aid cell-to-cell trafficking of movement-deficient mutants of turnip mosaic virus [20]. Thus, movement is potentially a highly evolvable redundant function of multiple genes that may support inter-cellular movement for the no-V2 monopartite begomoviruses. Alternatively, it was proposed that phloem-limited begomoviruses may have a reduced need for protein- mediated inter-cellular movement [27, 128], although this needs experimental assessment. The phloem-limited monopartite Americas begomoviruses ToLDeV [60] and ToMoLCV [27] provide an excellent system to explore this hypothesis. However, careful assessment of inter-cellular movement capabilities for these highly multifunctional proteins must be done on a case-by-case basis, given that (i) begomovirus positional homologs are not necessarily functional homologs [129] and (ii) many begomovirus proteins are also suppressors of RNA silencing and defective infection phenotypes may result from failure to suppress host defenses.

An additional feature of the V2/AV2-less Americas viruses is that their overall genomes/DNA-A are, on average, 130 nucleotides shorter than those from AAEO (Fig. 5). Ho et al. [26] attributed shorter segment lengths to progressive deletion of the V2/AV2 promoter region within the conserved region. We did not observe a systematic difference between CP and Rep lengths in our analyses, indicating that the difference likely lies in the length of the long intergenic region. As AAEO begomoviruses are thought to be ancestral to those in the Americas – in part because of the larger sequence diversity found in Africa, Asia, Europe and Oceania compared to begomoviruses in the Americas for both DNA-A proteins (this study) and DNA-B segments [54] – this implies a directionality to the difference: American begomoviruses likely experienced a genome reduction. Simplification and reduction in genome size may be the dominant mode of genomic evolution, occasionally interrupted by periods of complexification [130]. Additionally, high degree of overlapping gene regions in geminivirus genomes may suggest strong pervasive selection for smaller genomic segments [131, 132]. Interestingly, these shorter Americas DNA-As are accompanied by DNA-B segments that are, on average, 113 nt shorter than their AAEO counterparts [26].

### Sweepovirus CPs group outside other begomovirus CPs

Sweepovirus CP sequences display significant divergence from the rest of all the begomovirus CPs (Fig 2 and Fig S1). However, sweepovirus Reps are comfortably nested within the AAEO Rep sequences. The incongruencies between sweepovirus CP and Rep sequences suggest that they may be the product of recombination between a typical AAEO begomovirus for the Rep and a highly diverged begomovirus for the CP, or otherwise experienced selection pressures that drove the evolution of a unique CP through repeated fixation of mutations. Sweepoviruses are often the sister clade to all other begomoviruses in phylogenetic analyses [8, 10] and could resemble early begomoviruses more than other extant species. Of course, recombination among geminivirus genera may have created this relationship as well [6, 133–135]. One factor that could indicate sweepoviruses retain some ancestral begomovirus traits is their length, on average longer than the non-sweepovirus begomoviruses (Fig. 5). All of the closely related genera to *Begomovirus* have members with longer genomes than begomoviruses (≥2.9kb): *Maldovirus* [136], *Topocuvirus* [134, 136], *Grablovirus* [134, 136], *Turncurtovirus* [134, 136], *Opunvirus* [136], *Curtovirus* [134, 136], *Citlodavirus* [134, 136, 137], *Mulcrilevirus* [134, 136, 137]. Additionally, Fontenele et al. [138] speculate that a larger, citlodavirus-like ancestor may have been an intermediate step in the evolution of bipartite begomoviruses from a monopartite ancestor because citlodaviruses have a movement protein that resembles the one on begomovirus DNA-B segments.

### No evidence that CoYVV and CoGMV are ancestral to Americas-type begomoviruses

It was suggested that the genomic features of the CoYVV and CoGMV exemplars from Vietnam and basal relationship to Americas-type viruses for their complete DNA-A segments is evidence that Americas-type begomoviruses were present in Afro-Eurasia prior to continental separation [14, 74]. However, phylogenetic analyses revealed that the CP of CoYVV is closely related to the CP of contemporary Americas begomoviruses while the CoGMV CP is basal but also closely related to the Americas CP clade (Fig. 2). On the other hand, both the Rep of CoYVV and CoGMV cluster within the AAEO group (Fig. 3). These results suggest that CoYVV and potentially CoGMV may be relatively recent recombinants that inherited their CP from Americas begomoviruses and their Rep from AAEO begomoviruses, rather than exemplars providing evidence that Americas begomoviruses evolved from AAEO-type ancestors prior to the separation of the continents. The presented alternative that begomoviruses may have evolved in the AAEO region, and a progenitor of the current Americas begomoviruses was introduced more recently through human migration appears much more likely [8, 59]. Alternatively, the ancestry of the Americas CP sequences could involve recombination between a CoGMV-like virus and a begomovirus of unknown ancestry. Nonetheless, the CoYVV and CoGMV exemplars do not provide enough evidence to infer an origin for the Americas begomovirus clade. We recommend careful interpretation of DNA-A segment phylogenies since intermediate evolutionary relationships like the ones observed for the *Corchorus* viruses from Vietnam may be the product of recombination, as traditional phylogenetic analyses will average across distinct evolutionary histories within recombinant segments.

### Moving away from “New World/Old World” terminology

The Americas No-V2/AV2 and the AAEO groups are typically referred to as “New World” and “Old World” begomoviruses, respectively. In this work we have deliberately avoided these vaguely defined historical terms, which allude to the European rediscovery of the Americas in 1492. While there is a clear geographical pattern indicating that the AAEO and Americas lineages independently co-diversified with their hosts for a long period of time, three begomovirus species lacking V2/AV2 have been isolated from AAEO: from *Corchorus* species in Vietnam and India that often group with viruses from the Americas [14, 59, 139] and from soybean in Nigeria [61]. Despite current usage, we suggest that it does not make sense to use the phrase “New World” while referring to any of these three viral species. Further, all sweepoviruses can be considered more closely related to viruses from the AAEO region than the Americas, regardless of country of isolation, which may be related to the distinctive history of sweetpotato movement across the Pacific [140]. Finally, usage of the term “New World” is inconsistent outside of begomovirology. Many authors classify Australia as part of “the New World”, i.e., not part of Europe. Because the first sequenced isolate of tomato leaf curl virus was from Australia [141] and since that sequence clusters with sequences from African and Asia, we believe it is currently useful to consider Oceania together with Afro-Eurasia in defining begomovirus groups. (Note that the word “Oceania” is also used differently in different contexts).

Regardless, geographically named clades may be of decreasing importance in begomovirology in upcoming decades. Several important plant host-virus pairs have resulted from ‘new encounters’ [77, 81–83], and the two large geographically defined lineages are no longer fully distinct, as evidenced by the recombinant origin of tomato latent virus [57] after the worldwide spread of TYLCV.

## ACKNOWLEDGEMENTS

We would like to thank members of the Duffy lab at Rutgers University and our colleagues at the Hanley-Bowdoin lab in North Carolina State University for feedback and a critical reading of this manuscript. This work was supported by US NSF award OIA-1545553 to SD, an HHMI Gilliam Fellowship for Advanced Study for ACB. YBA was supported by the RISE program at Rutgers University.

---

**Figure S1.**
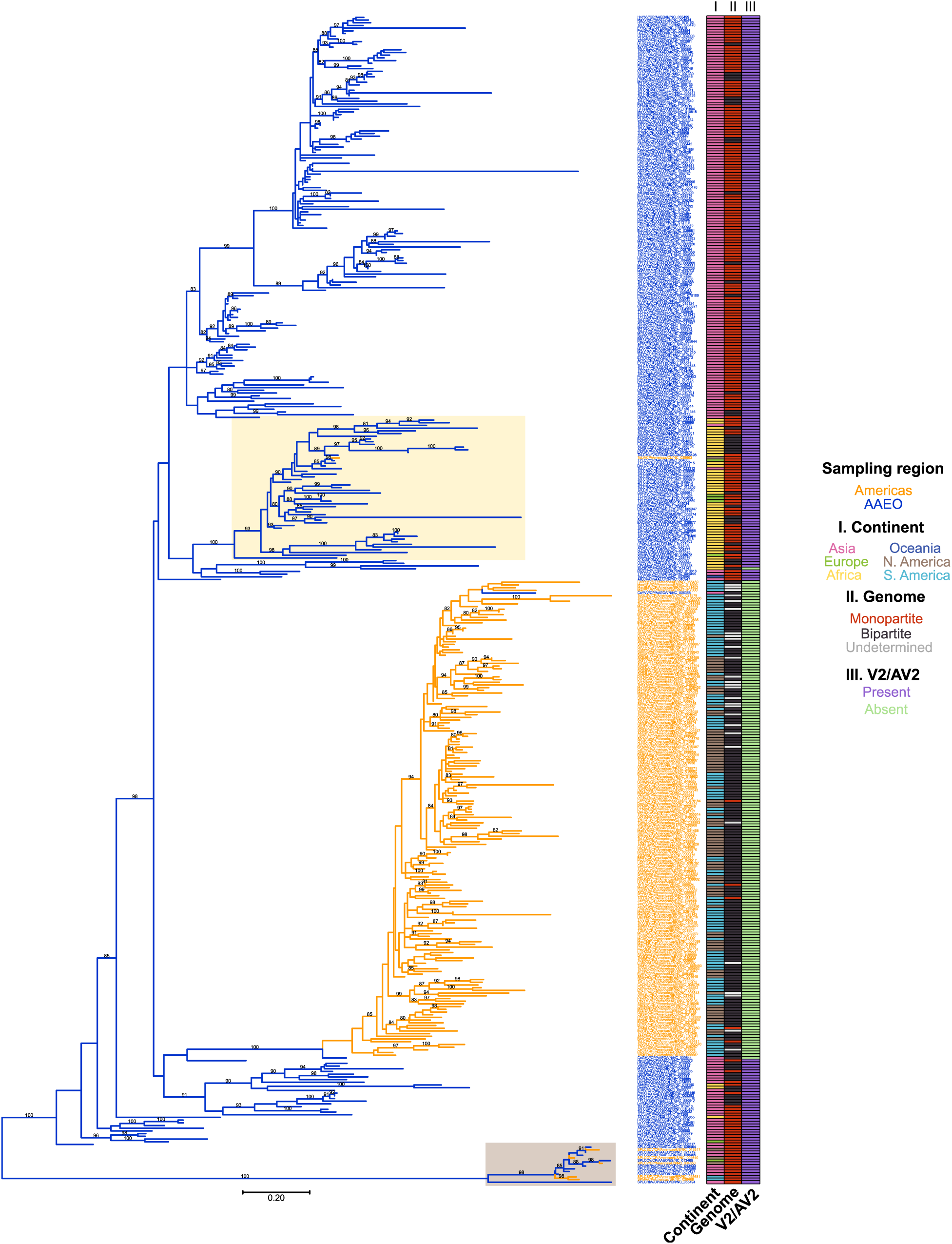
Midpoint-rooted maximum likelihood phylogenetic tree of complete amino acid sequences of 432 begomovirus RefSeq species exemplars. The maximum likelihood phylogenetic tree was constructed using IQ-Tree v2.07 with automatic selection of the best-fit substitution model (JTT+I+G4). Tree inference was performed with 3000 ultrafast bootstrap (UFBoot) replicates and a stopping rule of 500 iterations between unsuccessful improvements to the local optimum. UFboot branch support values ≥ 80% are shown mid-branch. The scale bar represents the number of substitutions per site. Branches are colored based on the region where the exemplar was sampled - Americas exemplars in orange and AAEO exemplars in blue. Sweepoviruses(brown) and the African clade (yellow) are highlighted. Exemplar labels follow the following format: “virus abbreviation/Rep/[Americas/AAEO]/two-letter country code/RefSeq accession number”.

**Figure S2.**
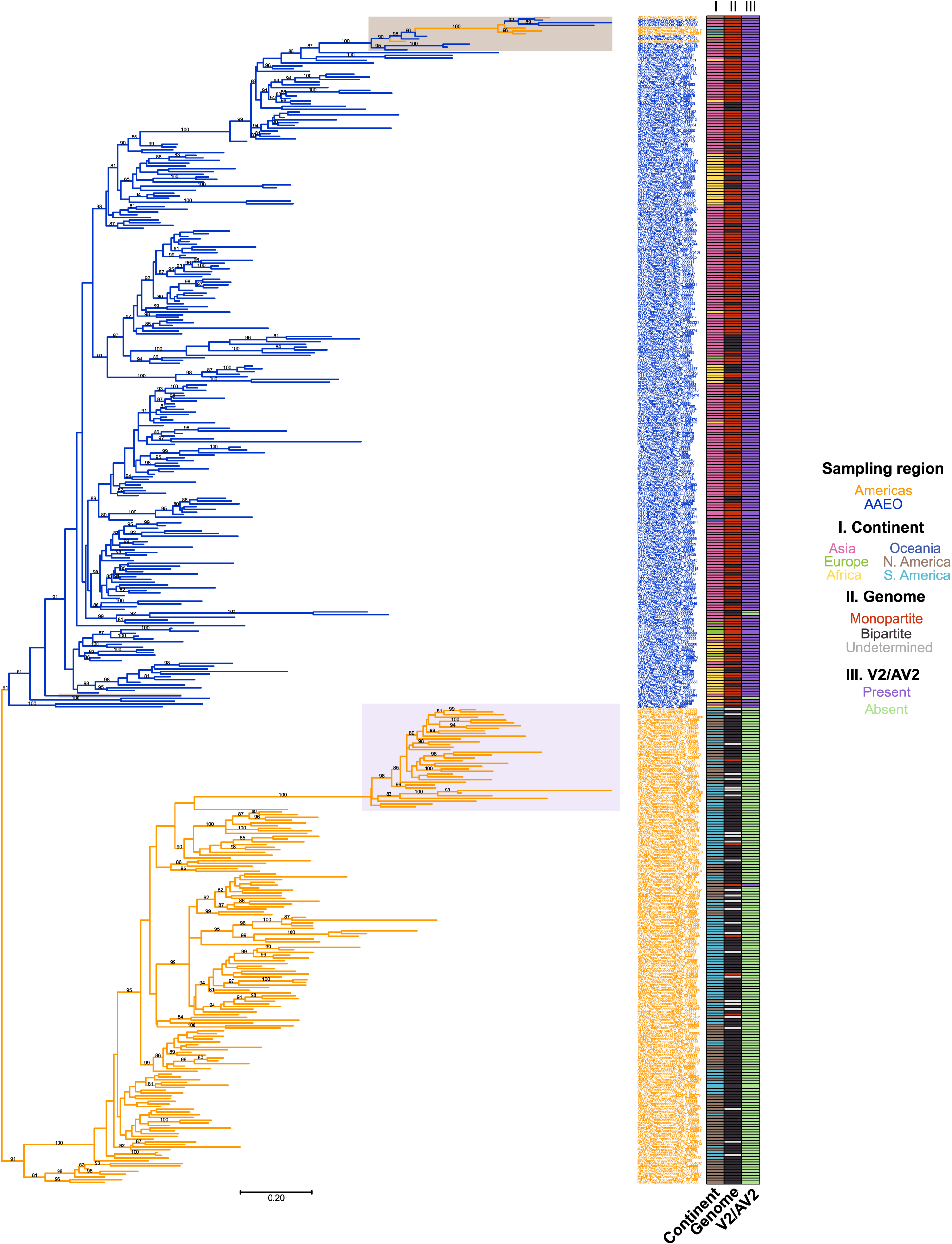
Midpoint-rooted maximum likelihood phylogenetic tree of trimmed Rep amino acid sequences of 432 begomovirus RefSeq species exemplars. The maximum likelihood phylogenetic tree was constructed using IQ-Tree v2.07 with automatic selection of the best-fit substitution model (LG+I+G4). Tree inference was performed with 3000 ultrafast bootstrap (UFBoot) replicates and a stopping rule of 500 iterations between unsuccessful improvements to the local optimum. UFboot branch support values ≥ 80% are shown mid-branch. The scale bar represents the number of substitutions per site. Branches are colored based on the region where the exemplar was sampled – Americas exemplars in orange and AAEO exemplars in blue. Sweepoviruses (brown) and Rep S-Lin clade (light red) are highlighted. Exemplar labels follow the following format: “virus abbreviation/Rep/[Americas/AAEO]/two-letter country code/RefSeq accession number”.

**Table S1.** Begomovirus species exemplars listed in the VMR that were not used in this study.

| VMR exemplars not used in this study (RefSeq accession number) | Reason |
| --- | --- |
| Bean golden yellow mosaic virus (NC_038791) | Truncated Rep |
| Chino del tomate Amazonas virus (NC_038443) | Significantly divergent Rep |
| Chilli leaf curl Bhavanisagar virus (NC_055130) | Truncated Rep |
| Cleome golden mosaic virus (NC_015397) | Significantly divergent and truncated Rep |
| Corchorus yellow vein mosaic virus (NC_020473) | Truncated Rep |
| Polygala garcinii virus (NC_037068) | Significantly divergent Rep |
| Sunn hemp leaf distortion virus (NC_013019) | Both CP and Rep listed as nonfunctional in GenBank (sequences not annotated as a result) |
| Sidastrum golden leaf spot virus (NC_038462) | Significantly divergent CP |
| Sida golden yellow spot virus (NC_038992) | Significantly divergent CP |
| Tomato golden leaf distortion virus (NC_043122) | Significantly divergent Rep |
| Tomato leaf curl Joydebpur virus (NC_074895) | Truncated CP and Rep |
| Tomato leaf curl Moheli virus (NC_038897) | Truncated Rep |
| West African Asystasia virus 3 | Does not have a RefSeq accession |

